# Intra-islet glucagon signalling regulates pulsatile insulin secretion and glucose homeostasis

**DOI:** 10.1101/2022.09.01.506223

**Authors:** K Suba, Y Patel, A Martin-Alonso, A Roberts, B Hansen, M Norton, J Shrewsbury, R Kwok, V Kalogianni, S Chen, X Liu, GA Rutter, B Jones, J Minnion, BM Owen, W Distaso, DJ Drucker, TM Tan, SR Bloom, KG Murphy, V Salem

## Abstract

**Background:** Type 2 diabetes (T2D) is characterised by the loss of pulsatile insulin secretion. We studied mice with β-cell specific loss of the glucagon receptor (Gcgr ^*fl/fl*^ X Ins-1^*Cre*^), to investigate the role of intra-islet glucagon receptor signalling on pan-islet calcium oscillations and insulin pulsatility.

**Methods:** Frequently sampled intravenous glucose tolerance tests were conducted on Gcgr ^β-cell-/-^ and littermate controls. Crossing with GCaMP6f (STOP flox) animals further allowed for β-cell specific expression of a fluorescent calcium indicator. These islets were functionally imaged *in vitro* and *in vivo*. Wild-type mice were transplanted with islets expressing GCaMP6f in β-cells into the anterior eye chamber and placed on a high fat diet. Part of the cohort received a glucagon analogue (GCG-analogue) for 40 days and the control group were fed to achieve weight matching. Calcium imaging was performed regularly during the development of hyperglycaemia and in response to GCG-analogue treatment.

**Results:** Gcgr ^β-cell-/-^ mice exhibited impaired glucose tolerance following intraperitoneal glucose challenge (control 12.7mmol/L ±0.6 vs. Gcgr ^β-cell-/-^ 15.4mmol/L ±0.0 at 15 min, p=0.002); fasting glycaemia was not different to controls. *In vitro*, Gcgr ^β-cell-/-^ islets showed profound loss of synchronised calcium waves in response to glucose which was only partially rescued *in vivo*. First-phase insulin pulsatility on peripheral blood sampling (n=5) was significantly disordered in Gcgr ^β-cell-/-^ mice (burst mass Gcgr ^β-cell-/-^ 0.30 ±0.03 versus 0.84 ±0.23 for controls p=0.04). Diet induced obesity and hyperglycaemia resulted in a loss of co-ordinated [Ca^2+^]_I_ waves in transplanted islets. This was reversed with GCG-analogue treatment, independently of weight-loss (n=8).

**Conclusion:** These data provide novel evidence for the role of intra-islet GCGR signalling in sustaining synchronised calcium oscillations and support a possible therapeutic role for glucagonergic agents to restore the insulin pulsatility lost in T2D.

## Introduction

The earliest hallmark of Type 2 diabetes (T2D) is the loss of pulsatile insulin secretion (1), which is thought to be central to the normal action of the hormone (2,3). Insulin pulsatility is closely related to islet-wide [Ca^2+^]_I_ oscillatory activity (4) and there is increasing attention turned towards understanding the islet of Langerhans as a functional unit. β-cell heterogeneity, specifically subpopulation of β-cell hubs or first responders (5,6), are thought to be important in the control of coordinated insulin release from pancreatic islets. Electrical coupling alone, via ionic gap junctions, is not enough to explain the observed connectivity patterns between β-cells (7), which prompted us to look at the modulatory role of paracrine factors, in particular α-cell-derived glucagon.

Glucagon is typically described as a counter-regulatory hormone to insulin, acting in the fasting state to raise blood glucose via hepatic glycogenolysis and gluconeogenesis. However, glucagon signalling has pleiotropic effects which are increasingly recognised, and its role in diabetes remains to be fully elucidated (8). The insulinotropic effects of glucagon have been known for many years (9). Glucagon infusions in healthy (10) and non-obese subjects with T2D (11) result in a rapid and substantial increase in plasma insulin levels under hyperglycaemic conditions. With the recent focus on targeting the glucagon receptor to treat diabetes, the role of intra-islet glucagon signalling is of increasing interest. To date, most of our understanding of the intra-islet effects of glucagon has come from *in vitro* or *ex vivo* studies (12,13,14) using isolated islets and high concentrations of glucagon. These corroborate the insulinotropic effects of glucagon, but attribute them, at least in part, to β-cell GLP-1 receptor activation (15,16,17,18), since intra-islet levels of glucagon are likely much higher than systemic levels and glucagon can activate the GLP-1 receptor at high concentrations (19). Overall, it remains poorly understood how intra-islet glucagon augments β-cell function, over what dynamic range, and the physiological and pharmacological relevance of intra-islet glucagon receptor activation.

We aimed to characterise the effects of β-cell specific loss of the glucagon receptor (GCGR) on islet connectivity and pulsatile insulin secretion. We hypothesised that intra-islet GCGR signalling contributes to the maintenance of islet-wide β-cell functional connectivity and pulsatile insulin secretion. We then examined whether glucagon receptor agonist therapy in the setting of obesity-induced dysglycaemia directly restores islet β-cell functional connectivity.

## Results

### GcgrKO^*β-cell* -/-^ islets appear histologically normal and show normal expression levels of β-cell identity and “disallowed” genes

Mice on a C57bl6/J background were bred to knock out the glucagon receptor (GCGR) specifically in β-cell (Gcg ^*β-cell*-/-^) using the model of Gcgr^*fl/fl*^ crossed with an Ins1^*Cre*^ line (see Methods for full details). The islets of these mice were isolated at 12 weeks of age. Quantification of mouse Gcgr mRNA using qPCR on whole islets revealed a large reduction in Gcgr expression in Gcgr ^*β-cell* -/-^ animals versus littermate controls (Figure 1A. Relative expression 0.55 ±0.04 for control vs 0.14 ±0.10 for Gcgr ^*β-cell*-/-^; *p=0.02*). There was no evidence of compensatory upregulation of GLP-1 or GIP receptor expression levels in Gcgr ^*β-cels*-/-^ islets (Supplementary Figure 1A and B). The anatomy of *Gcgr*^*β-cell* -/-^ islets was unperturbed (Figure 1B), and there were no differences in relative β-cell mass (β- to α-cell ratio) when compared with littermate controls (Supplementary Figures 1C-E). mRNA expression of established β-cell identity markers (*Pdx1, Mafa, Kcnj11*) and “disallowed” (*Acot7, Ldha*) genes (20) were also comparable to control levels (Figure 1C). Taken together, these experiments suggest that germline Ins1^*Cre*^-driven deletion of GCGR in β-cells is not associated with changes in islet maturation or architecture.

**Figure 1.**
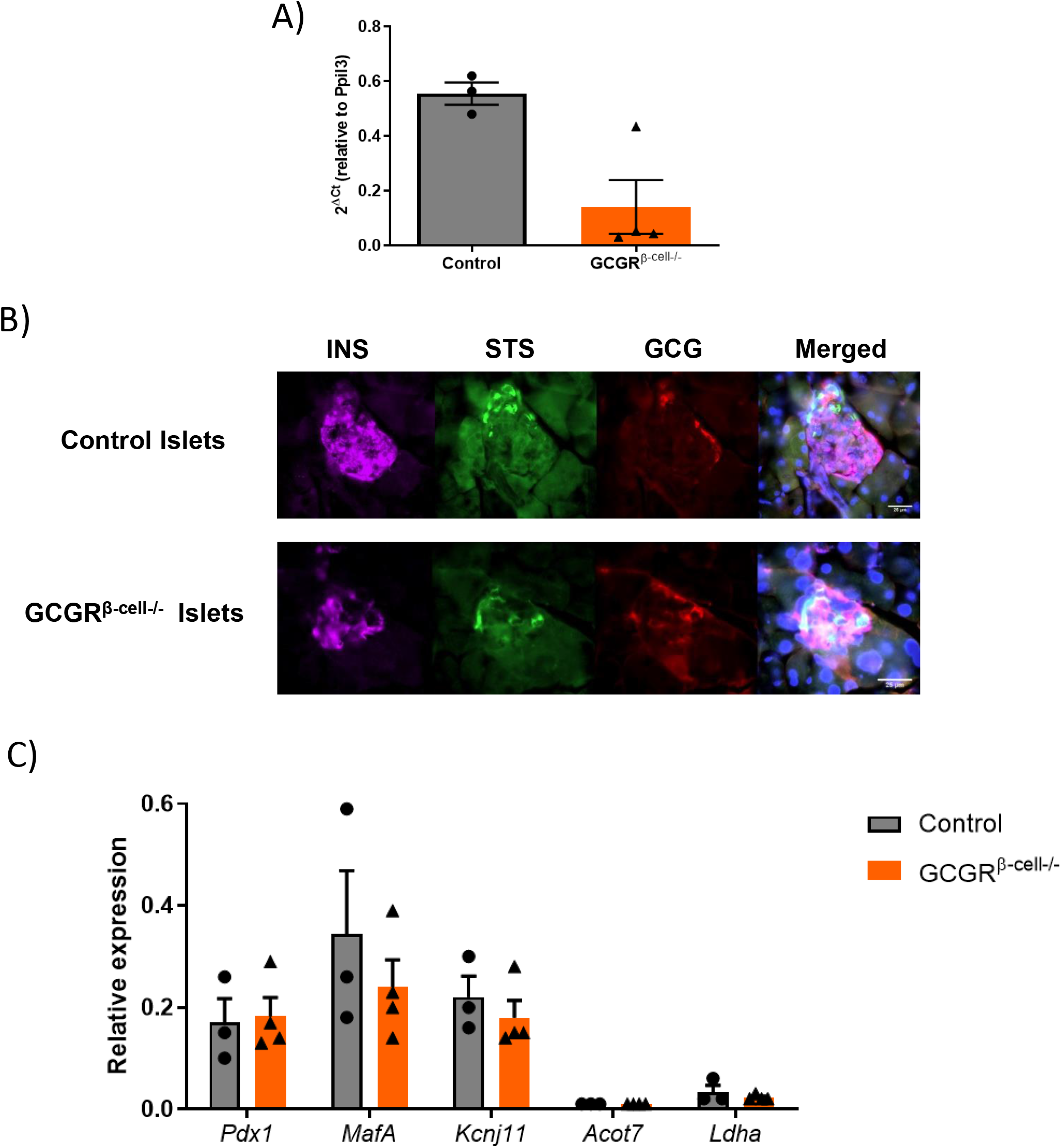
β-cell specific deletion of the glucagon receptor does not result in morphological changes to islets or β-cells. A) The expression of Gcgr relative to *Ppil3* is reduced from 0.55 in littermate control islets to 0.14 in Gcgr ^*β-cell*-/-^ islets (unpaired t-test n=4 animals per genotype, p=0.0187). B) Control and Gcgr ^*β-cell*-/-^ pancreata (n=2 per genotype) were stained for insulin, somatostatin and glucagon. C) Relative expression of β-cell identity genes *Pdx1, Mafa, Kcnj11, Acot7* and *Ldha* is not altered in Gcgr ^*β-cell*-/-^ islets (n=3) versus controls (n=4; unpaired t-tests; p=ns).

### Gcgr^*β-cell* -/-^ mice exhibit an insulin secretory deficit

Unlike mice with a whole body deletion of the glucagon receptor (21), our model of β-cell specific loss of the glucagon receptor did not result in a body weight phenotype: adult body weight at 12 weeks was 31.7 ± 0.9 g for controls (n=10) versus 29.9 ±0.9 g for Gcgr ^*β-cell*-/-^ animals (n=15) (*p=ns*; Figure 2A). Circulating glucose levels following five hours fasting were similar in Gcgr ^*β-cell*-/-^mice and controls, but during an intraperitoneal glucose tolerance test (IPGTT), circulating glucose concentrations were significantly higher at 15 min. and 30 min. following glucose in Gcgr ^*β-cell*-/-^ animals (Figure 2B-C). Insulin tolerance (ITT) and hepatic glucose mobilisation (as measured with a pyruvate tolerance test) were not different in Gcgr ^*β-cell*-/-^ animals (Supplementary Figures 2A and B), and basal levels of circulating insulin, glucagon and GLP-1 were also similar to controls (Supplementary Figures 3A-D). In this context, the reduced glucose tolerance suggested impaired glucose-stimulated insulin secretion (GSIS) in Gcgr ^*β-cell*-/-^ animals. To further test this, we conducted hyperglycaemic clamp experiments on Gcgr ^*β-cell*-/-^ animals and littermate controls (Figure 2D). We found that the glucose infusion rate (GIR) was significantly lower in the (weight-matched) Gcgr ^*β-cell*-/-^ cohort (66 mg/kg/min ± 0.3 versus 83 mg/kg/min ±0.1 for controls; *p=0.001*). Taken together, these experiments suggest that β-cell loss of GCGR signalling results in impaired early GSIS.

**Figure 2.**
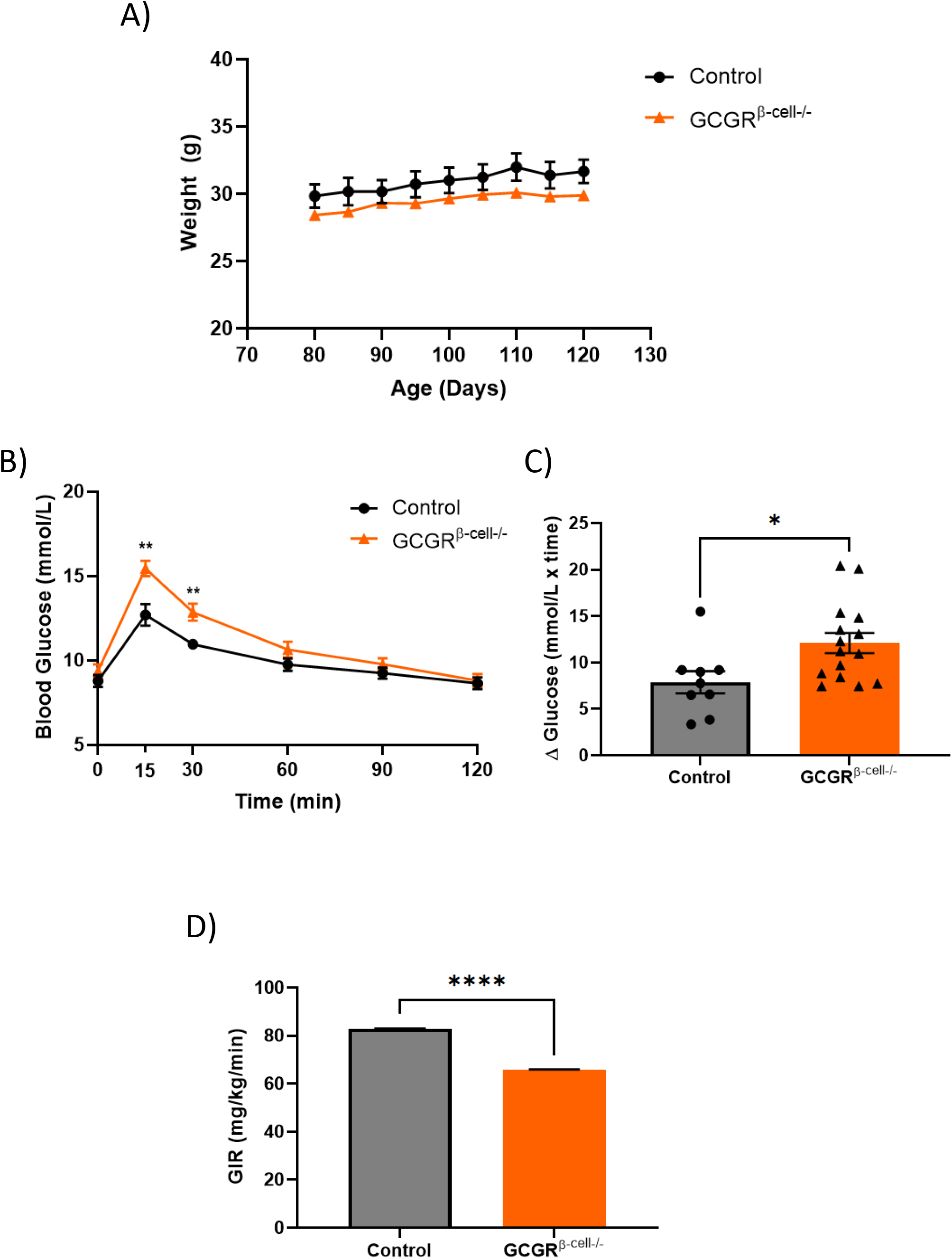
*Gcgr*^*β-cell*-/-^ mice are glucose intolerant. A) The body weight of 12-week old Gcgr ^*β-cell*-/-^ animals (n=15) is comparable to littermate controls (n=10). B) Gcgr ^*β-cell*-/-^ animals show significantly increased blood glucose concentrations at 15min. and 30min. time-points in an IPGTT (control vs Gcgr ^*β-cell*-/-^, t=15, 12.7mmol/L ±0.6, 15.4mmol/L ±0.5; t=30, 11.0mmol/L ±0.3, 12.9mmol/L ±0.5; control n=9; Gcgr ^*β-cell*-/-^ n=15; Two-tailed, Unpaired t-test; p=0.002). C) AUC measurements of whole glucose excursion were also significantly higher in Gcgr ^*β-cell*-/-^ animals during the IPGTT (Two-tailed, Unpaired t-test; p=0.02). D) Hyperglycaemic (blood glucose: 16.5mmol/L ±1.5) clamp experiments revealed a lower glucose infusion rate in Gcgr ^*β-cell*-/-^ animals compared with age-matched controls (66mg/kg/min ±0.3 versus 83mg/kg/min ±0.1 for GCGR^*β-cell*-/-^ versus control animals, respectively; Gcgr ^*β-cell*-/-^ n=5, control n=2; Two-tailed, Unpaired t-test on AUC; p<0.001).

### Pulsatile insulin secretion is altered in Gcgr^*β-cell* -/-^ mice

To further characterise first-phase insulin secretory pattern of Gcgr ^*β-cell* -/-^ animals, we frequently sampled circulating insulin levels following an intravenous (IV) glucose bolus (Figure 3A). This revealed that the average burst mass of insulin pulses is significantly increased in Gcgr ^*β-cell* -/-^ animals (0.30 ± 0.03 versus 0.84 ± 0.23 for controls and Gcgr ^*β-cell*-/-^respectively) in response to a glucose challenge. However, burst mass was variable in Gcgr ^*β-cell*-/-^animals but highly consistent in the control group (Figure 3B). After an early burst, there was a tendency for fewer subsequent insulin peaks in the Gcgr ^*β-cell*-/-^animals. The average duration of a pulse showed a tendency to be longer in the Gcgr ^*β-cell* -/-^ (2.9 ± 0.4 versus 3.8 ± 0.6 mins for control versus Gcgr ^*β-cell*-/-^) mice and to occur at a reduced frequency (2.9 ± 0.4 versus 3.8 ± 0.6 mins for control versus Gcgr *^β-cell-/-^; p=ns*). Together these findings suggest that pulsatile insulin secretion is altered in the absence of intra-islet GCGR signalling and this may contribute to the insulin secretory deficit observed previously.

**Figure 3.**
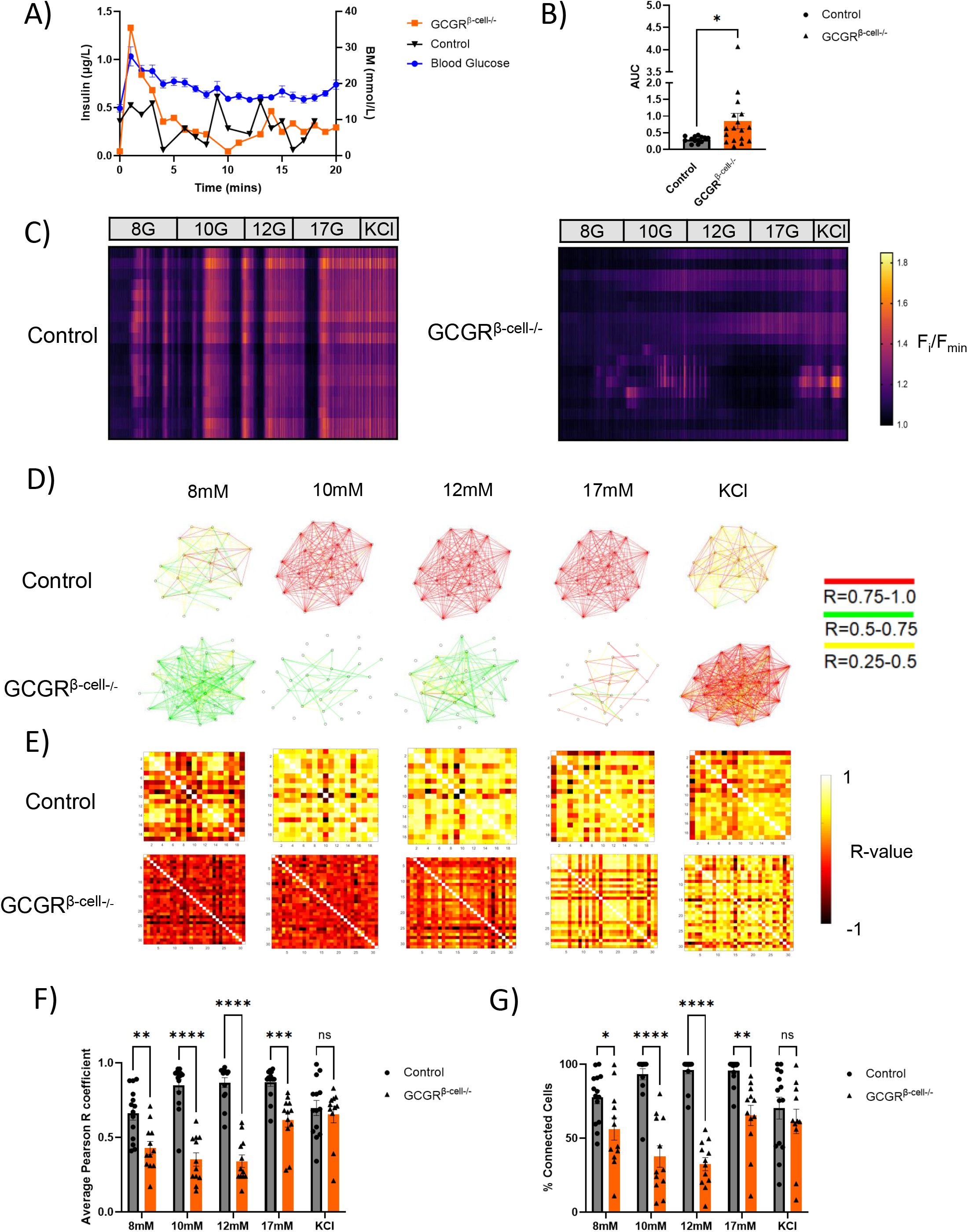
Pulsatile insulin secretion and *in vitro* islet [Ca^2+^]_I_ dynamics are altered in Gcgr ^*β-cell*-/-^ animals. A) Representative blood glucose and insulin curves from control and Gcgr ^*β-cell*-/-^ animals during a frequently sampled intravenous glucose tolerance test (FSIVGTT). B) Average burst mass during an FSIVGTT is significantly increased in Gcgr ^*β-cell*-/-^ animals compared to age-matched controls (control n=4, Gcgr ^*β-cell*-/-^ n=5; 0.30 ±0.03 versus 0.84 ±0.23 for controls and Gcgr ^*β-cell*-/-^respectively, Two-tailed, Mann-Whitney test; p=0.03). C) Isolated islets from controls and Gcgr ^*β-cell*-/-^ islets that also expressed the calcium fluorophore GCaMP6f in their β-cells were imaged in a perifusion system with rising glucose concentrations terminating with the addition of KCl. These raster plots show mean intensity changes in individual islet segments (18 segments per islet) during the imaging session. D) Cartesian connectivity maps and E) heat maps of Pearson R values between β-cells of control and Ins1^*Cre*^GCaMP6f^*fl/fl*^: Gcgr ^*β-cell*-/-^ islets *in vitro* at 8-17mM glucose concentrations and following KCl treatment. F) The average Pearson co-efficient in control animals is significantly higher than in Ins1^*Cre*^GCaMP6f^*fl/fl*^: Gcgr ^*β-cell*-/-^ animals across all glucose conditions (Two way ANOVA, Mann-Whitney test; **p=0.001 and ***p<0.0001).G) The average percentage connectivity was significantly elevated at all glucose concentrations in control islets compared to Ins1^*Cre*^GCaMP6f^*fl/fl*^: Gcgr ^*β-cell*-/-^islets (n=14 islets from 3 animals for control group and n=12 islets from 3 animals for Ins1^*Cre*^GCaMP6f^*fl/fl*^: Gcgr ^*β-cell*-/-^ group; Two way ANOVA, Mann-Whitney test; *p=0.04 and ***p<0.0001).

### *In vitro* calcium imaging reveals that β-cell GCGR loss affects coordinated [Ca^2+^]_I_ activity

Islet (β-cell) intracellular calcium [Ca^2+^]_I_ oscillations underpin insulin secretion (22). To better understand the mechanisms driving the insulin secretory deficit in Gcgr ^*β-cell*-/-^ mice, these animals were further crossed with a line expressing the [Ca^2+^]_I_ indicator GCaMP6f selectively in the β-cells. *In vitro*, Ins1^*Cre*^GCaMP6f^*fl/fl*^: Gcgr ^*β-cell*-/-^ and control (Ins1^*Cre*^GCaMP6f^*fl/fl*^: Gcgr ^*β-cell* +/+^) islets were exposed to step-wise increases in glucose concentration on a perifusion stage. Segmental analysis of optical section-wide [Ca^2+^]_i_ activity revealed that at 8mM glucose concentration and above, synchronised [Ca^2+^]_I_ oscillations, spanning the entire islet cross-section, were evident in all control islets (n=13 islets from 3 animals; Supplementary Video 1; Figure 3C left Raster plot). On the other hand, no global increase in oscillatory [Ca^2+^]_I_ activity was evident in any of the Ins1^*Cre*^GCaMP6f^*fl/fl*^: Gcgr ^*β-cell*-/-^ islets studied (n=18 islets from 4 animals; Figure 3C right Raster plot). Instead, localised rises in [Ca^2+^]_I_ were observable (Supplementary Video 2), which passed our threshold of Δ20% Fi/Fmin for recording an increase in [Ca^2+^]_I_, suggesting that β-cells were responding in a stochastic manner but not producing coordinated, islet-wide pulses. The addition of 40mM KCl to Gcgr ^*β-cell*-/-^ islets elicited a strong pan-islet [Ca^2+^]_I_ response, suggesting that a powerful depolarising stimulus is able to overcome the defective communication between β-cells (Figure 3D-G). Taken together, these observations suggest that β-cell GCGR signalling is important for the generation of synchronised [Ca^2+^]_I_ responses to glucose *in vitro*. Defective β-cell communication and the lack of synchronisation may partially explain the insulin secretory deficit noted in Gcgr ^*β-cell*-/-^ animals.

### *In vivo* [Ca^2+^]_I_ imaging reveals that the loss of co-ordinated calcium activity in Gcgr^*β-cell*-/-^ islets is only partially rescued *in vivo*

*In vitro* perifusion studies cannot recapitulate the complex microenvironment of *in situ* islets, where their activity may be modulated by neuronal input and a capillary bed. To directly investigate β-cell connectivity of Gcgr ^*β-cell*-/-^ in vivo, Ins1^*Cre*^GCaMP6f^*fl/fl*^: Gcgr ^*β-cell*-/-^-expressing islets were implanted into the eyes of syngeneic WT recipients, where they act as faithful “reporters” that can be directly imaged (23) (control: n=8 islets in 3 recipients; Gcgr ^*β*-cell-/-^: n=7 islets in 3 recipients; Figure 4A). Following full implantation, islet [Ca^2+^]_I_ dynamics were recorded under high circulating glucose (10.4 ± 1.5 mmol/L and 11.3 ± 0.4 mmol/L for Ins1^*Cre*^GCaMP6f^*fl/fl*^: Gcgr ^*β-cell*-/-^ and control groups respectively; *p=ns*). In the islets from control animals, 7 out of 8 islets were observed to have cross-sectional calcium wave activity as opposed to only 4 out of 7 of the Gcgr ^*β*-cell -/-^ islets. We analysed single cell [Ca^2+^]_i_ dynamics in the oscillating islets, applying statistical methods to measure correlations between β-cell [Ca^2+^]_I_ activity, using previously described methods (6). The average Pearson R values of Ins1^*Cre*^GCaMP6f^*fl/fl*^: Gcgr ^*β*-cell-/-^ islets was significantly lower than in the control islets (R=0.88 ± 0.06 compared to R=0.98 ± 0.02, p=0.036) (Figure 4B). In parallel, the average proportion of connected cells was non-significantly lower in islets lacking β-cell Gcgr (82.1 ±12.2% compared to 97.1 ± 1.96% for control islets; *p=ns*) (Figure 4C). We conclude that the detrimental effects of Gcgr knockout on β-cell [Ca^2+^]_I_ dynamics are only partially rescued post implantation *in vivo*.

**Figure 4.**
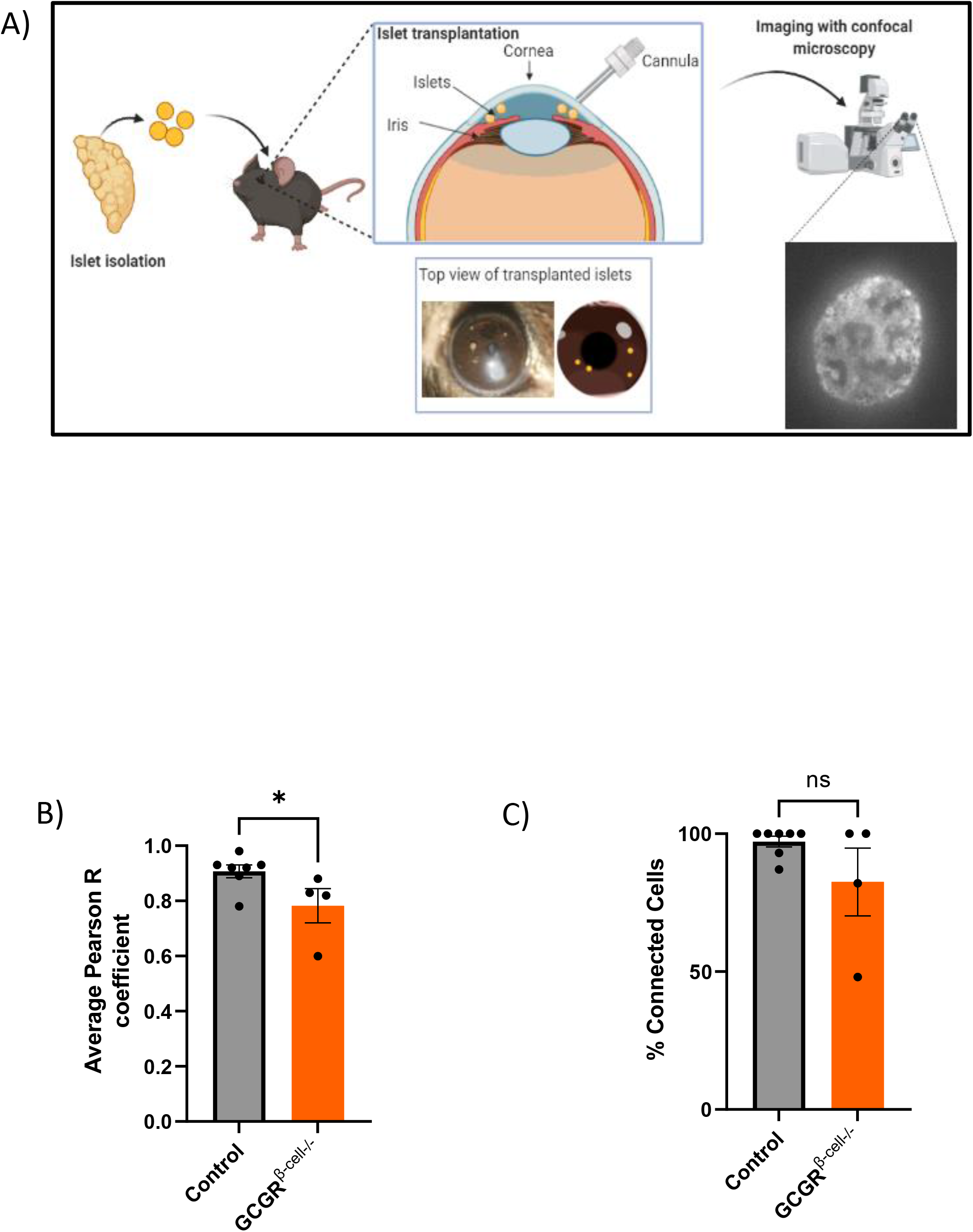
Partial restoration of β-cell connectivity in GCGR^*β-cell*-/-^ islets following implantation in the murine eye. A) Schematic of the experimental imaging platform. Islets with β-cell specific loss of the GCGR that also express the [Ca^2+^]_I_ (Ins1^*Cre*^GCaMP6f^*fl/fl*^: Gcgr ^*β-cell*-/-^-islets) are isolated and transplanted into the anterior camber of the eye of a syngeneic recipient. Ins1^*Cre*^GCaMP6f ^*β-cell*+/+^-islets were implanted as a control. After 4 weeks, the islets implant, are innervated and functional. Thereafter they can be directly imaged (confocal microscopy) longitudinally B) A total of 7 Ins1^*Cre*^GCaMP6f^*fl/fl*^: Gcgr ^*β-cell*-/-^-islets and 7 Ins1^*Cre*^GCaMP6f^*β-cell*+/+^-islets were imaged under high circulating glucose conditions (in three animals). Six control islets and only four Ins1^*Cre*^GCaMP6f^*fl/fl*^: Gcgr ^*β-cell*-/-^ islets exhibited [Ca^2+^]_I_ waves. Using single cell connectivity readouts as described in Figure 3, we assessed these [Ca^2+^]_I_ waves for β-cell connectivity. The average coefficient of connectivity was significantly lower in Ins1^*Cre*^GCaMP6f^*fl/fl*^: Gcgr ^*β-cell*-/-^ islets (Two-tailed, Mann-Whitney test; p=0.036) and C) the average number of connected β-cells was lower but this effect was not significant (Two-tailed, Mann-Whitney test; p=ns).

### Diet-induced obesity results in reduced β-cell connectivity which can be restored by chronic administration of a GCG-analogue

Selective deletion of the Gcgr in β-cells revealed a possible beneficial physiological role for GCGR signalling in islet functional connectivity. Consequently we investigated whether pharmacological GCGR agonism in a diet-induced obese (DIO) model of chronic hyperglycaemia can alleviate the deleterious effects of high-fat diet on co-ordinated insulin secretion (Figure 5A). We created a glucagon analogue (GCG-analogue, see Methods and Supplementary Figure 4 for sequence) with strong potency for cAMP production at the mouse glucagon receptor (m Gcgr) but not different than native glucagon at the mGLP-1R. The analogue was 164 times more potent at the mouse glucagon compared to the mouse GLP-1 receptor (m Gcgr >mGLP-1R: ΔΔpEC50 = 2.2 ± 0.3).

**Figure 5.**
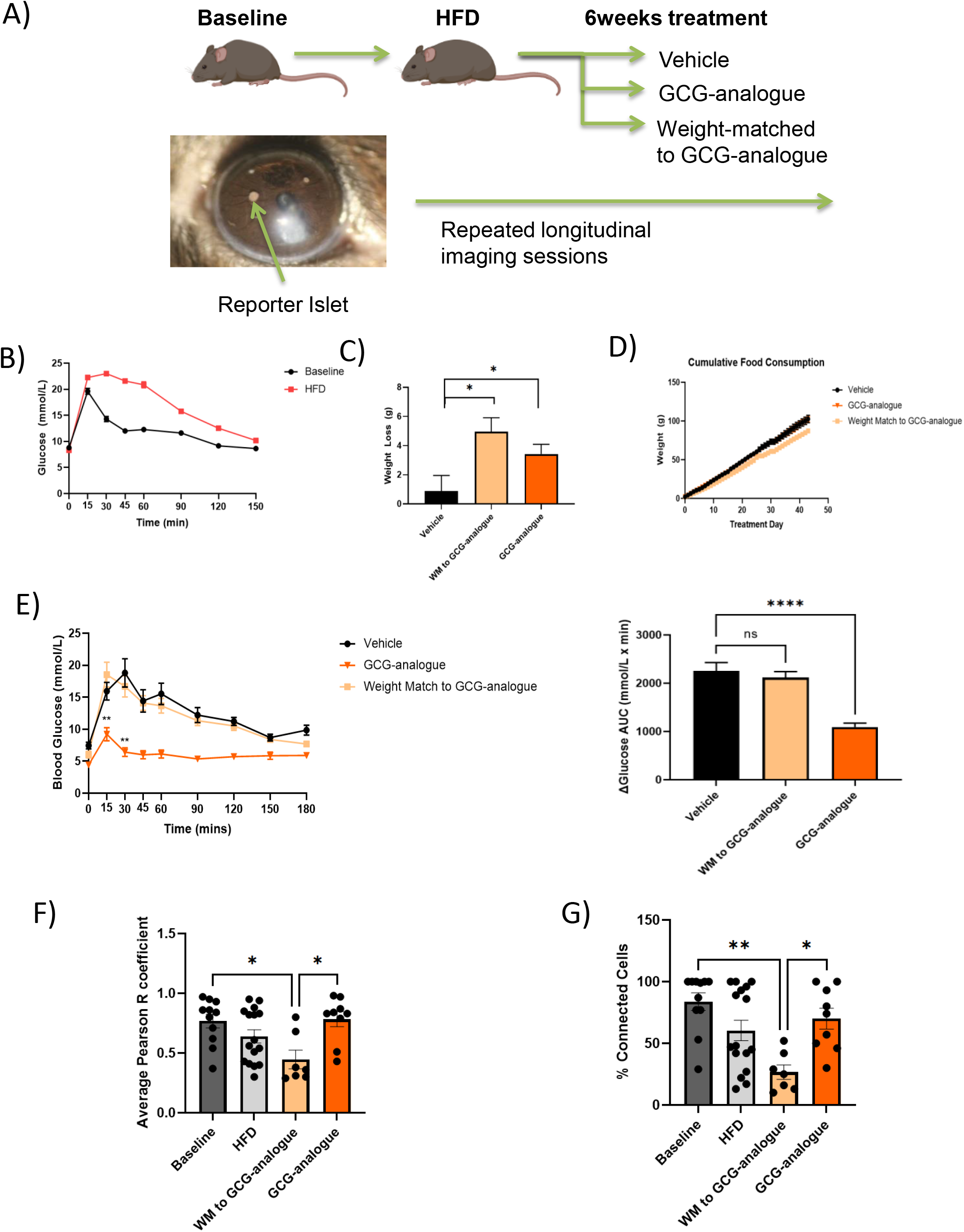
Direct observation of islet function following the induction of diet-induced obesity in mice and subsequent treatment with a synthetic glucagon-analogue (GCG-analogue). A) Schematic diagram of the study. B) High-fat feeding of 10-week old mice leads to the development of diet-induced obesity and impaired glucose tolerance (n=63-59; the AUC at baseline was 508.3 ±106 versus 1309 ±154.9 after HFD; Two-tailed Welch’s t-test; p<0.0001). C) The mean bodyweight was 39.6 ±0.9g prior to the start of the treatment and was comparable across groups. GCG-analogue treatment caused a significant weight-loss of −12.01% ±1.97 versus −2.3% ±2.7 in the vehicle group (n=8 per group treatment group, one-way ANOVA, Bonferroni multiple comparison test; p=0.029). D) Cumulative food consumption over the course was comparable between the three treatment groups (one-way ANOVA, Tukey’s test; p=ns). E) GCG-analogue treatment improves glycaemia of DIO mice. Glucose concentrations at the 15min. and 30min. time-points of an IPGTT were significantly lower in the GCG-analogue group compared to the weight-matched group (t15=9.2mmol/L ±1 versus 12.5mmol/L ±2 for GCG-analogue versus WM to GCG-analogue groups p=0.001; t30=6.4mmol/L ±0.7 versus 16.7mmol/L ±1.6 for GCG-analogue versus WM to GCG-analogue groups p=0.001; One-way ANOVA; Tukey’s test). F) HFD results in reduced β-cell connectivity and this is rescued by the GCG-analogue. The average Pearson Co-efficient is significantly increased in the GCG-analogue group compared with the weight-matched group (Kruskal-Wallis test; p=0.015) and significantly lower in the weight-matched group compared to baseline (Kruskal-Wallis test; p=0.015). G) The average percentage connectivity is significantly higher in the GCG-analogue group than in the weight-matched group at the end of 40 days treatment (Baseline n=11 islets; HFD n=16 islets; WM to GCG-analogue n=7 islets; GCG-analogue n=9 islets; Kruskal-Wallis test; p=0.045).

Twenty-four lean C57BL6J mice (aged 8 weeks) received syngeneic transplants of Ins1^*Cre*^GCaMP6f^*fl/fl*^-expressing “reporter” islets into their anterior eye chamber and were then placed on a high fat diet (HFD). Following 12 weeks of the HFD, the mice demonstrated diet-induced obesity (DIO) and impaired glucose tolerance (Figure 5B). Animals were then randomised by body weight into three groups of 8: Group 1 received a daily dose of GCG-analogue, Group 2 were control/vehicle-injected and fed *ad libitum* and Group 3 received daily control/vehicle injections but were food restricted to weight match (WM) to Group 1. Forty days of GCG-analogue treatment resulted in a body weight loss in Group 1 of 12.01% ±1.97 (*p=0.030*; Figure 5C). The cumulative food intake in Group 3 was significantly less than both Groups 1 and 2, supporting previous reports of an increased energy expenditure mechanism for weight loss with exogenous administration of a glucagon agonist (24) (Figure 5D).

Treatment with the GCG analogue produced a weight loss independent normalisation of glucose homeostasis in DIO animals (Figure 5E). Fasting glucose was lower (4.3mmol/L versus 7.5mmol/L; *p=0.005*) and intraperitoneal glucose tolerance was (peak glucose of 9.2mmol/L versus 18.5mmol/L in GCG-analogue versus weight matched-groups, respectively; *p<0.001*).

In order to understand the direct islet effects of GCG-analogue, we monitored the activity of the reporter islets in these animals across the duration of the study. Firstly we demonstrate, for the first time longitudinally and *in vivo*, a disruption in β-cell coordinated [Ca^2+^]_I_ activity related to obesity. The worsening glycaemic phenotype of DIO mice was paralleled by a diminution of pan-islet β-cell connectivity that was recovered by GCG-analogue treatment but not weight loss alone (Figures 5F and G). Of note, blood glucose levels throughout the imaging sessions were comparable (10.6 ±0.52 mmol/L versus 12.7±0.6 mmol/L versus 10±0.3 mmol/L over three imaging sessions), therefore differences in [Ca^2+^]_I_ responsiveness were unlikely to have been a consequence of low circulating glucose levels during the imaging studies. Taken together, our findings indicate that DIO impairs coordinated [Ca^2+^]_I_ responses of β-cells and that these coordinated responses can be restored by GCG-analogue treatment via mechanisms that are independent of weight loss.

## Discussion

This study investigated the physiological role of intra-islet glucagon signalling and the pharmacological effects of glucagon receptor activation in islets. We found that β-cell specific deletion of the Gcgr results in a glucose intolerant phenotype that is redolent of early T2D, with a loss of normal, pulsatile first phase insulin secretion. We also provide evidence of the utility of GCGR-preferring peptide analogues for the recovery of obesity-induced dysglycaemia.

Our model of β-cell specific GCGR deletion resulted in a large reduction in whole-islet measured Gcgr transcript, consistent with reports that most of the GCGR in the islet resides on β-cells (25). The glucose intolerant phenotype in these mice was associated with an insulin secretory deficit without any changes to islet architecture or β-cell mass. Furthermore, the intra-islet receptor expression and serum levels of two major incretins GLP-1 and GIP, were not different from controls, suggesting a lack of compensatory morphological or functional changes that are common in global enteropancreatic-hormone receptor knockout models. Zhang et al. recently reported a glucose intolerant phenotype in their model of β-cell GCGR loss although they did not report on pulsatility (26). Conversely, others who have studied intra-islet GCGR signalling in germline (27) or conditional (14) β-cell knock out of the GCGR using a mouse insulin promoter (MIP)^*Cre*^-driven mechanism have not reported glucose intolerance. This may be due to differences in promoter activity. *Ins1* promoter-driven systems, and particularly the Ins-1^*Cre*^ knock-in model used here, are highly specific to β-cells (29, 30), with no reported extra-pancreatic expression or alterations to glucose homeostasis, whilst some groups have reported that an ectopic expression of the human growth hormone (hGH) minigene as well as an increased β-cell mass and insulin content (31) occur with the MIP promoter. Therefore, differences in the animal models used to investigate islet paracrinology must be taken into account when comparing studies.

We propose that the highly specific loss of the GCGR on β-cells in mice results in a phenotype that resembles some of the earliest manifestations of T2D. The pulsatile release of insulin has long been recognised as an important feature of glucose regulation – discouraging insulin-receptor downregulation. There is strong evidence that pulsatile delivery of insulin to the liver is necessary to maintain normal insulin receptor activity (2) and it has been proposed that the loss of insulin pulsatility *per se* is the cause of hepatic insulin resistance (3). Loss of insulin pulsatility occurs at an even earlier stage than the frank diminution of first phase insulin secretion during the transition from impaired glucose tolerance to T2D, and it is also seen in the normoglycaemic first-degree relatives of patients with T2D (1). Fascinating work by Nunemaker et al. revealed that the shape of the individual secretory bursts from individual islets and from *in vivo* sampling are almost superimposable (corrected for amplitude), suggesting that the *in vivo* pulse is generated by simultaneous secretion from an imprinted islet population (4). Therefore, understanding the islet-level structure/function relationships that sub-serve insulin pulsatility is important.

*In vitro*, Gcgr ^*β-cell*-/-^ islets possessed β-cells that were able to respond in a stochastic manner to rises in glucose but failed to mount any co-ordinated waves of activity over the range of 3 to 17mM glucose. This supports a paracrine role of glucagon in maintaining pan-islet β-cell functional connectivity. A recent study found *in vitro* evidence for the critical importance of glucagon in the first-phase response of islets to a glucose stimulus. The first-phase GSIS response is conditional on cAMP signalling which, above a certain threshold, plays a permissive role for insulin secretion. Thus the paracrine effect of glucagon may be important for maintaining β-cell cAMP-tone (32). Intriguingly, glucagonergic stimulation of islets has been shown to drive oscillatory cAMP activity (33, 34) which synchronises with [Ca^2+^]_I_ responses in β-cells. When transplanted into the anterior eye chamber of the mouse, islets become vascularised and receive innervation patterns homologous to those in the pancreas (35,36). Around a half of the Gcgr ^*β-cell*-/-^ islets transplanted in the eye did not mount secretory wave activity, and in those that did, measures of β-cell connectivity were significantly reduced. In the *in vivo* environment the cAMP tone of β-cells may be supported by a multitude of other signals. For example, cholinergic receptor activation has also been shown to have synchronising properties in islets (37). Deciphering which are the most important factors will help the development of novel treatments that restore co-ordinated insulin release. Given the differences in the topographical relationships of α- and β-cells across species, as well as differences in aspects of innervation, it will be important to investigate the relevance of these findings to human islets.

To better understand the impact of metabolic insults on β-cell [Ca^2+^]I oscillations *in vivo*, we functionally assessed reporter islets longitudinally in the eye of DIO animals with a hyperglycaemic phenotype. The induction of DIO caused a diminution in pan-islet [Ca^2+^]_I_ waves and β-cell connectivity read-outs, suggesting that high-fat feeding interferes with the ability of β-cells to propagate [Ca^2+^]_I_ across the islet. Whilst some have questioned the potential decoupling of insulin secretory activity from [Ca^2+^]_I_ dynamics with the use of certain anaesthetic agents (including isoflurane which was used in these experiments), we note that these *in vivo* findings mirror numerous previous *ex vivo* and *in vitro* reports of the deleterious effects of lipotoxicity on coordinated insulin secretory behaviour (38, 39). The precise mechanism by which HFD causes a loss of synchronised oscillations is presently unknown. However, it has been postulated that the deleterious effects of FFAs include suppressing the expression of gap junctions and reducing islet responsiveness to incretins (39).

In recent years, dual GLP-1R/GCGR agonists have gained increasing research attention due to their potential for superior weight-lowering effects and improved glycaemic control compared with GLP-1-based monotherapies (40, 11). As an example, cotadutide is currently being tested in Phase 2 clinical trials following its promising effects on DIO mice and non-human primates (41). The additional weight loss evoked by cotadutide is attributed to GCGR-driven mechanisms to increase energy expenditure. Concurrently, GLP-1R agonism is thought to counterbalance any glucose-raising aspect of the glucagonergic element and to improve the glycaemic profile of DIO mice. However, after adjusting for weight loss, cotadutide-treated animals still show better glucose response curves than liraglutide-treated DIO mice (42). We sought to investigate the direct effects of glucagon receptor activation in a clinically relevant model of obesity-associated hyperglycaemia, using a peptide agonist which was over ten times more potent at the glucagon than the GLP-1 receptor. Chronic treatment with this GCGR agonist for 40 days resulted in a markedly improved glycaemic profile in DIO mice. This improvement in glucose homeostasis coincided with a 12 ±2% weight loss, yet the restoration of β-cell connectivity and co-ordinated pan islet [Ca^2+^]I activity measured with the GCGR-analogue was not recapitulated with equivalent weight loss alone, supporting the assertion that GCGR-activation promotes β-cell connectivity independently of weight loss. This raises the possibility that GCGR agonism may be particularly useful in restoring insulin secretory function in patients who are unable to lose enough pancreatic fat with dieting to resolve their T2D.

## Future Directions and Conclusion

Pancreatic islets are micro-organs which integrate numerous neural, vascular and paracrine regulatory inputs to fine-tune their hormone secretory output to maintain glucose homeostasis. In this study, we provide evidence for the role of intra-islet GCGR signalling in coordinated [Ca^2+^]_I_ oscillations and first-phase insulin release *in vivo*. The phenotype of Gcgr ^*β-cell*-/-^ animals is altered first-phase insulin secretory responses during an intravenous glucose challenge, redolent of pre-diabetes. Future studies should aim to fully elucidate the contributions of endocrine and paracrine signalling, gut hormone receptor distribution and function, to co-ordinated islet activity. This is necessary to guide future multi-agonist development for the treatment of T2D.

## ACKNOWLEDGEMENTS

VS designed and supervised the study, with thanks to KGM, GAR, SRB, TT and DD for guidance. KS, YP, AMA, AR, MN, and VS undertook the mouse studies. VS, WD and BH developed scripts and KS, JS, RK, VK, SC and SL contributed to connectivity analyses. SRB designed the glucagon analogue which was developed and tested by BJJ and JM. VS, KGM and KS wrote the manuscript contributions from all authors.

VS is supported by a Diabetes UK Harry Keen Clinician Scientist Fellowship (15/0005317). G.A.R. was supported by a Wellcome Trust Investigator Award (212625/Z/18/Z), MRC Programme grant (MR/R022259/1) a start-up grant from the CHUM Research Center at the University of Montreal and a Canadian Foundation for Innovation John R. Evans infrastructure grant. DJD was supported by CIHR grant 154321, a Banting and Best Diabetes Centre Novo Nordisk Chair in Incretin Biology, and the Sinai Health Novo Nordisk Foundation fund in regulatory peptides.

## Supplementary Figure Legends

**Supplementary Figure 1.**
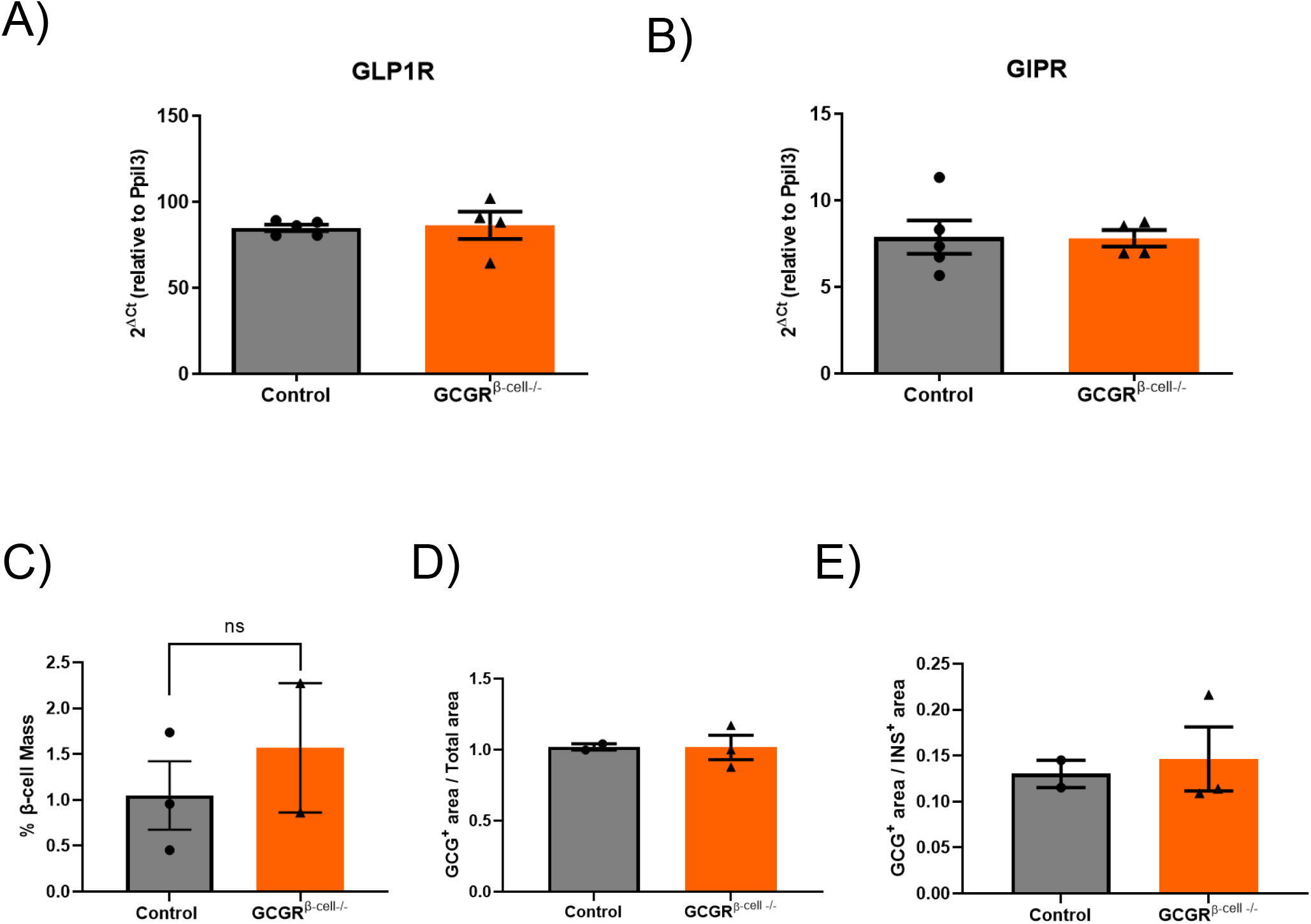
GLP-1 and GIP receptor expression and the relative α- and β-cell content is unaltered in the islets of Gcgr ^*β-cell*-/-^ animals. A) Relative GLP-1R expression levels were 84.9(±1.9) for controls versus 86.4 (±7.9) for Gcgr ^*β-cell*-/-^ animals (Two-tailed, Mann-Whitney test; p=ns). B) Relative GIPR expression levels were 7.8(±1) for controls versus 7.8 (±0.5) for Gcgr ^*β-cell*-/-^ animals (Two-tailed, Mann-Whitney test; p=ns). C) β-cell mass relative to total pancreatic tissue area is not significantly different in Gcgr ^*β-cell*-/-^ (n=2) versus control (n=3) animals (Two-tailed, Mann-Whitney test; p=ns). D) α-cell mass relative to total pancreatic tissue area and E) α- to β-cell ratio is not significantly different in Gcgr ^*β-cell*-/-^ (n=3) versus control animals (n=2; Two-tailed, Mann-Whitney test; p=ns).

**Supplementary Figure 2.**
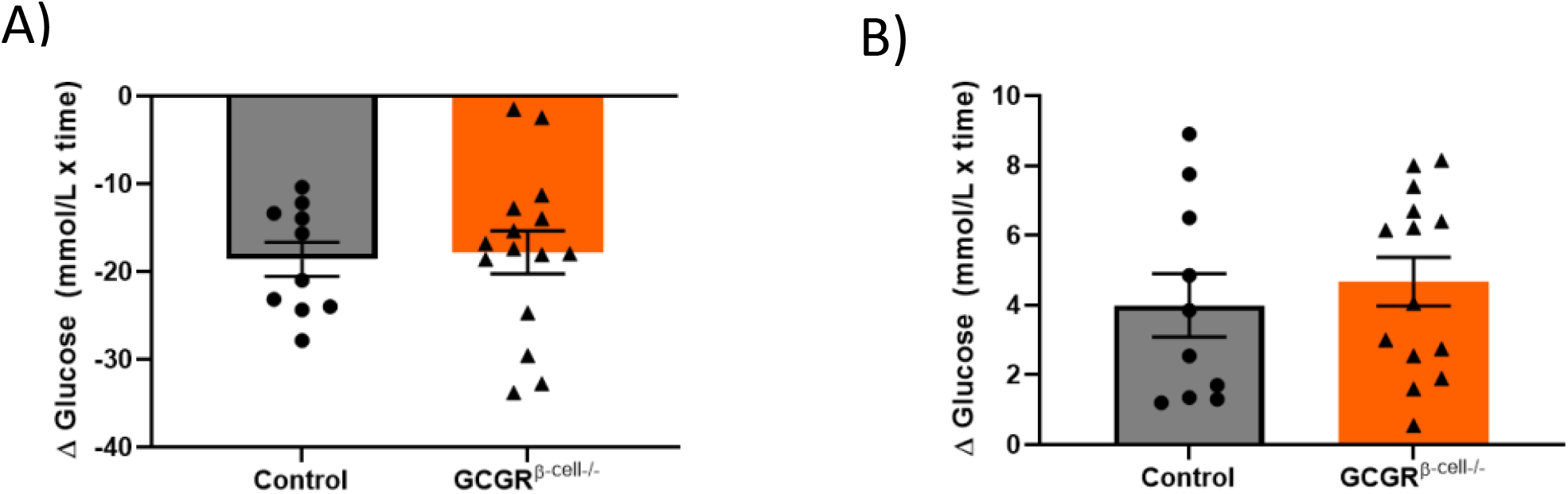
There is no difference in glucose excursion following insulin or pyruvate tolerance tests between Gcgr^*β-cell*-/-^ and littermate control animals. Insulin (A) and pyruvate (B) tolerance testing is unaltered in Gcgr ^*β-cell*-/-^ animals (n=15) compared to controls (n=10; Two-tailed, Unpaired t-tests and Mann-Whitney test; p=ns).

**Supplementary Figure 3.**
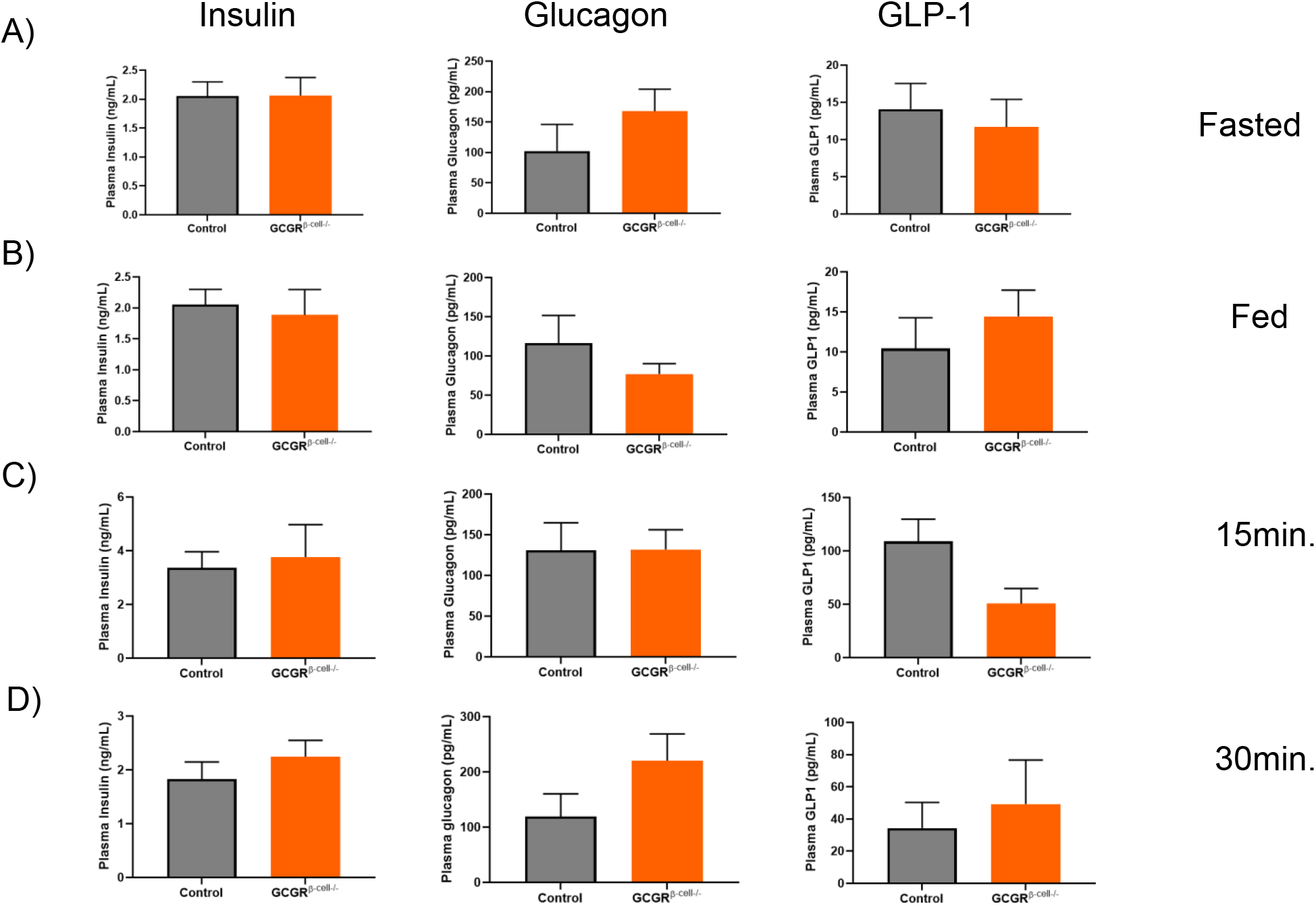
Serum hormone levels during an IPGTT are comparable Gcgr^*β-cell*-/-^ and control animals. Glucagon and GLP-1 levels in A) fasted, B) non-fasted, C) at 15min. and D) 30min. during an IPGTT are comparable between control and Gcgr^*β-cell*-/-^ animals (Two-tailed, Unpaired t-test of timed measurements; p=ns). Whilst insulin pulsatility is affected (see Figure 3) absolute levels of insulin at timepoints in group GTT tests were not significantly different.

**Supplementary Figure 4.**
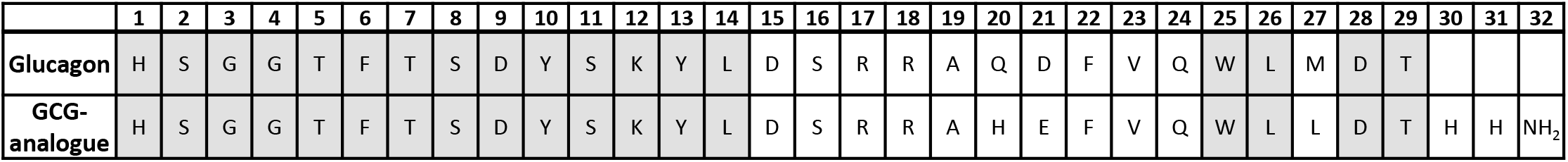
The synthetic GCG-analogue. A synthetic analogue of native glucagon was generated with amino acid changes in positions 21 (from aspartate to glutamate) and 27 (from methionine to leucine) and with a further addition of two histamine groups preceding an amine group.

## Methods

### Animals

All procedures involving animals were conducted in accordance with the UK Animal (Scientific Procedures) Act 1986, under Project Licence PPL 75/0462. Gcgr ^*fl/fl*^ mice on C57BL6 background were generated by inserting a locus of X-over P1 (*LoxP*) site upstream of exon 6 on chromosome 11. A flippase recognition target (*FRT*) and *LoxP*-flanked neomycin resistance cassette was inserted downstream of exon 12. Animals with the neomycin resistance cassette were crossed with mice expressing flippase (*FLP*) to remove the *FRT*-insert, resulting in litters with *LoxP* sites at exons 6 and 12 of chromosome 11. For the β-cell specific deletion of the GCGR, Gcgr^*fl/fl*^ animals were crossed with Ins1^Cre^-expressing animals (42). The Ins1^*Cre*^GCGR^*fl/fl*^ mouse line was bred in-house and produced normal litter sizes. Mice expressing the Ins1^*Cre*^ construct and homozygous for Gcgr ^*fl/fl*^ are referred to as ‘Gcgr ^*β-cell*-/-^’ within the text. Mice that do not express *Cre recombinase* under the Ins1 promoter (Ins1^*Cre*−^ GCGR^*fl/fl*^) were used as controls and referred to as ‘littermate controls’ within the text.

For [Ca^2+^]_I_ imaging studies, Ins1^*Cre*^-expressing mice were crossed with mice that expressed GCaMP6f^*fl/fl*^ fluorescent calcium sensor, downstream of a *LoxP*-flanked STOP cassette (The Jackson Laboratory, stock no. 028865) and bred in-house. For calcium imaging studies with GCGR^*β-cell*-/-^ islets, the Ins1^*Cre*^GCGR^*fl/fl*^ mouse line was crossed with Ins1^*Cre−*^ GCaMP6f^*fl/fl*^ mice for β-cell specific GCGR deletion and expression of the calcium sensor GCaMP6f. These mice are referred to as ‘GCaMP6f: Gcgr ^*β-cell*-/-^’ within the text. Lastly, Ins1^*Cre*^GCaMP6f^*fl/fl*^ animals, with intact β-cell GCGR-signalling and the GCaMP6f calcium reporter, were used as controls to GCaMP6f: Gcgr ^*β-cell*-/-^ islets and in the chronic GCG-analogue study.

For experiments with the GCG-analogue, Ins1^*Cre*^GCaMP6f^*fl/fl*^-expressing islets were transplanted into the anterior eye chamber of C57BL6/J syngeneic wild-type (WT) recipients (Envigo, Huntingdon UK). All animals were maintained under controlled conditions (21-23°C; 12:12 hr light:dark schedule) and following full implantation (>4weeks) implanted islets (n=3-4 per animal) were imaged: at baseline (normal chow), two months after HFD (5.21kcal/g; 20% kcal protein, 60% kcal fat and 20% kcal carbohydrate; Research Diets, D12492) to induce DIO, and 40 days of daily sub-cutaneous injections of vehicle (saline+Zn^2+^; n=8) or GCG-analogue (n=8) and after weight-matching (n=8) intervention. The synthetic GCG-analogue was administered subcutaneously. Metabolic tests (intraperitoneal glucose and insulin tolerance tests) were conducted at baseline, after HFD (to ascertain DIO) and after treatment.

### Metabolic Testing

Intraperitoneal glucose tolerance tests (IPGTTs) (2g/kg), intraperitoneal insulin tolerance tests (IPTTs) (1U/kg) and intraperitoneal pyruvate tolerance tests (IPPTTs) (2g/kg) throughout this study were conducted after 5hrs fasting. Blood glucose measurements were taken at −15, 0, 15, 30, 60 and 120min time-points.

### Hyperglycaemic Clamp Experiments

Under isoflurane anaesthesia (2%), indwelling catheters were surgically implanted into left common carotid artery and right jugular vein of Gcgr ^*β-cell*-/-^ mice and littermate counterparts, and allowed to recover for 7 days. 50% glucose solution was continuously infused into the jugular vein while 10U/ml heparinised saline was inserted into the carotid artery to withdraw blood samples during a hyperglycaemic clamp. GIR was measured over 45 minutes of plateaued glucose levels (typically achieved at 20 minutes after clamp start).

### Frequently Sampled Intravenous Glucose Tolerance Test (FSIVGTT)

Following a 5hrs fast, Gcgr ^*β-cell*-/-^ mice and littermate controls received an indwelling catheter into their left common carotid artery, under isoflurane anaesthesia (2%) and were allowed to recover for 7 days. Blood samples (6μL) were taken for baseline blood glucose and insulin measurements. Animals received a bolus of *D*-glucose (50% solution; 100μL) and the peripheral blood was sampled for glucose and insulin measurements at 1 minute intervals thereafter, for up to 20minutes in total. The insulin concentration of blood samples were measured using the Mercodia mouse insulin ELISA kit, as per manufacturers’ instructions.

### qPCR Analysis

RNA from tissue samples was extracted using the guanidium thiocyanate-phenol-chloroform method. To quantify gene expression both probe-based (TaqMan^®^ gene expression assay) and dye-based (SYBR^®^ Green JumpStart^™^ Taq ReadyMix^™^ kit) methods were used. Reactions were performed in CFX384^™^ detection system (Bio-Rad).

### Immunohistochemistry

For immunohistochemistry (IHC) analysis, pancreata were sectioned at 10μm intervals and slices stained with antibodies for insulin (conjugated with Alexa Fluor^®^ 594 dye), somatostatin (conjugated with Alexa Fluor^®^ 488 dye), and glucagon (conjugated with Alexa Fluor^®^ 647 dye) and DAPI. Images were captured using Eclipse Ti2 inverted microscope (Nikon) with an ORCA-Flash 4.0LT+ digital camera (Hamamatsu, UK) and 40x objective.

### Quantification of IHC of Pancreas Sections

Quantification of IHC images of pancreatic slices were conducted blindly. Briefly, images were thresholded for each fluorophore to generate binary images of the islet fluorescence signal. Areas of increased fluorescence were measured over binary images.

### Insulin, Glucagon and GLP-1 Immunoassays

Hormones from blood samples of Gcgr ^*β-cell*-/-^ mice and littermate controls were measured using standard ELISA kits (for insulin and glucagon, 62IN3PEF and 62SGLPEB, Cisbio, France; GLP-1, 81508, Crystal Chem, UK).

### Peptide generation and receptor activation (cAMP) assay

The GCG-analogue analogue was designed and donated by Professor SR Bloom (Imperial College London). Peptides were synthesised by Bachem (UK) and underwent local testing for purity after aliquoting and freeze drying, according to well in the Bloom Drug Development programme. Cyclic AMP (cAMP) analysis was conducted using DiscoverX PathHunter cells expressing human GCGR, GLP-1R or GIPR using the cAMP Dynamic 2 cAMP kit (Cisbio, France) as per manufacturer’s instructions.

### *In vivo* imaging experiments

Islets implanted in the anterior eye chamber were imaged (ex.:488nm; at 3Hz) under isoflurane anaesthesia (<2%) using a spinning disk confocal microscope (Nikon Eclipse Ti, Crest spinning disk, 20x water dipping 1.0 NA objective) under high circulating glucose.

### *In vitro* perifusion experiments

*In vitro* perifusion experiments were recorded using Yokogawa CSU22 Nipkow spinning disk microscope coupled with a Zeiss Axiovert M200 and x10/0.3NA objective (Zeiss) and Volocity software. Islet [Ca^2+^]_I_ dynamics (ex.:488nm; exp. time: 500msec) were recorded at 1Hz. In *in vitro* perifusion studies, Ins1^*Cre*^GCaMP6f^*fl/fl*^: Gcgr ^*β-cell*-/-^-expressing and control (Ins1^*Cre*^GCaMP6f^*fl/fl*^ Gcgr ^*β-cell*+/+^ islets were exposed to step-wise increases in glucose concentration from 2mM to 12mM and 17mM for 8 minutes per condition. For glucagon analogue (GCG-analogue) experiments, Ins1^*Cre*^GCaMP6f ^*fl/fl*^ islets were exposed to 3mM, 7mM and 17mM glucose Krebs solutions for 8 minutes per condition. Islets were perfused with glucose-only Krebs solution (control) or 60nM GCG-analogue dissolved in Krebs solution.

### Statistical Analyses

Statistical significance between two conditions was assessed using the paired or unpaired, Student’s *t*-test. Interactions between multiple conditions were determined using one- or two-way analysis of variance (ANOVA) (with Tukey’s or Bonferroni’s post-hoc tests). Analyses were performed using GraphPad Prism (GraphPad Software v.8.0) and MATLAB (Mathworks) and significant *P* values are described in each relevant section. Values are plotted as mean±s.e.m., unless otherwise stated.

